# Deciphering the Sex Bias in Housekeeping Gene Expression in Adipose Tissue: a Comprehensive Meta-analysis of Transcriptomic Studies

**DOI:** 10.1101/2021.12.04.471124

**Authors:** Maria Guaita-Cespedes, Rubén Grillo-Risco, Marta R. Hidalgo, Sonia Fernández-Veledo, Borja Gómez-Cabañes, Gonzalo Anton-Bernat, Helena Gomez-Martinez, Deborah Jane Burks, María de la Iglesia-Vayá, Amparo Galán, Francisco Garcia-Garcia

**Affiliations:** Bioinformatics and Biostatistics Unit, Principe Felipe Research Center (CIPF), Valencia, 46012, Spain; Hematology Research Group, Instituto de Investigación Sanitaria La Fe (IIS La Fe), Valencia, 46026, Spain; Department of Endocrinology and Nutrition and Research Unit, University Hospital of Tarragona Joan XXIII, Institut d’Investigaciò Sanitària Pere Virgili (IISPV), Tarragona, Spain; CIBER de Diabetes y Enfermedades Metabólicas Asociadas (CIBERDEM), Instituto de Salud Carlos III, Madrid, Spain; Molecular Neuroendocrinology Laboratory, Principe Felipe Research Center (CIPF), Valencia, 46012, Spain; Imaging Unit FISABIO-CIPF, Fundación para el Fomento de la Investigación Sanitaria y Biomédica de la Comunidad Valenciana, 46012, Valencia, Spain

**Keywords:** housekeeping genes, meta-analysis, transcriptomics, sex bias, adipose tissue

## Abstract

**Background:** As the housekeeping genes (HKG) generally involved in maintaining essential cell functions are typically assumed to exhibit constant expression levels across cell types, they are commonly employed as internal controls in gene expression studies. Nevertheless, HKG may vary gene expression profile according to different variables introducing systematic errors into experimental results. Sex bias can indeed affect expression display, however, up to date, sex has not been typically considered as a biological variable.

**Methods:** In this study, we evaluate the expression profiles of six classical housekeeping genes (four metabolic: *GAPDH, HPRT, PPIA*, and *UBC*, and two ribosomal: *18S* and *RPL19*) to determine expression stability in adipose tissues (AT) of *Homo sapiens* and *Mus musculus* and check sex bias and their overall suitability as internal controls. We also assess the expression stability of all genes included in distinct whole-transcriptome microarrays available from the Gene Expression Omnibus database to identify sex-unbiased housekeeping genes (suHKG) suitable for use as internal controls. We perform a novel computational strategy based on meta-analysis techniques to identify any sexual dimorphisms in mRNA expression stability in AT and to properly validate potential candidates.

**Results:** Just above half of the considered studies informed properly about the sex of the human samples, however, not enough female mouse samples were found to be included in this analysis. We found differences in the HKG expression stability in humans between female and male samples, with females presenting greater instability. We propose a suHKG signature including experimentally validated classical HKG like *PPIA* and *RPL19* and novel potential markers for human AT and discarding others like the extensively used *18S* gene due to a sex-based variability display in adipose tissue. Orthologs have also been assayed and proposed for mouse WAT suHKG signature. All results generated during this study are readily available by accessing an open web resource (https://bioinfo.cipf.es/metafun-HKG) for consultation and reuse in further studies.

**Conclusions:** This sex-based research proves that certain classical housekeeping genes fail to function adequately as controls when analyzing human adipose tissue considering sex as a variable. We confirm *RPL19* and *PPIA* suitability as sex-unbiased human and mouse housekeeping genes derived from sex-specific expression profiles, and propose new ones such as *RPS8* and *UBB*.

**Highlights:**

- A computational strategy based on massive data analysis revealed that an accurate experimental design for adipose tissue requires the adequate selection of a sex-unbiased housekeeping genes (HKG).
- The extensively used 18S gene displays sex-based variability in adipose tissue, although PPIA and RPL19 do not, and hence, represent robust HKG.
- New sex-unbiased human and mouse candidate HKG: RPS8 and UBB.
- metafun-HKG (https://bioinfo.cipf.es/metafun-HKG): a freely available web tool to allow users to review stable expression levels of candidate HKG along the large volume of FAIR data.

## Introduction

Housekeeping genes (HKGs) are a large class of constitutively expressed genes subjected to low levels of regulation under various conditions. They generally perform biological actions fundamental to basic cellular functions such as the cell cycle, translation, metabolism of RNA, and cell transport [1,2]. Thus, the stable expression of HKGs is assumed in all cells of an organism independent of the tissue, developmental stage, cell cycle state, or presence/absence of external signals [3,4].

The use of internal controls when performing quantitative gene expression analysis (such as microarrays, RNA-sequencing [RNA-seq], and quantitative reverse transcriptase-polymerase chain reaction [qRT-PCR]) represents the most common strategy to normalize gene expression to correct for intrinsic errors related to sample manipulation and the technical protocol. The gene expression profiles obtained depend significantly on the reference genes employed as internal controls; therefore, inappropriate controls can lead to inaccurate results.

Given their fundamental roles, HKGs tend to display medium-high expression levels; this characteristic makes these genes especially suitable as internal controls/reference genes to normalize gene expression data in quantitative gene expression analysis [2,5,6]. Ideally, internal controls should exhibit stable gene expression across most sample types and experimental conditions to minimize undesired experimental variation; however, the literature suggests that the expression of commonly used HKGs varies depending on the experimental conditions and chosen setup and the analyzed tissue [5-13]. Importantly, these limitations do not invalidate the use of HKGs as a normalization strategy; instead, they support the need for a deeper understanding of how HKGs behave under different conditions or in distinct tissues. The stability of HKG expression must be validated under the particular conditions of interest of each study as a mandatory step [5], considering all experimental, biological, or clinical variables [7, 14-16]. Importantly, this should include sex as an essential variable.

The role of sex in biomedical studies has often been overlooked, despite evidence of sexually dimorphic effects in biological studies. Karp *et al*. recently demonstrated how sex phenotypically influenced a substantial proportion of mammalian traits, both in wildtype and mutants [17]. Meanwhile, Oliva *et al*. reported the impact of sex on gene expression in various human tissues through metadata analysis by the GTEx platform, generating a catalog of sex-based differences in gene expression and the regulatory pathways involved [18]. The authors revealed ubiquitous effects of sex on gene expression; however, they highlighted significant sex-based differences in human visceral and subcutaneous adipose tissue. Sex as an intrinsic variable has not been historically considered of immense importance. In a recent review of more than 600 animal research studies, 22% of publications did not specify animal sex [19]. Of the reports that specified animal sex, 80% of publications included only males and 17% only females, leaving only 3% that considered animals of both sex [20]. An analysis of the number of animal studies revealed a more significant disparity-16,152 males vs. only 3,173 females. Only seven studies (1%) reported sex-based results. Thus, the number of male-only studies and the use of male animals have become more disparate over time [20,21]. Unfortunately, human counterpart studies do not provide any encouragement; while international institutions now consider sex as a critical variable [22,23], the male perspective predominates in past studies. The lack of consideration of sex as a variable can accentuate/attenuate gene expression analysis, which has subsequent implications on biological or biomedical interpretations.

The quantitative analysis of gene expression data has allowed assessments of gene expression levels within different tissues and under various conditions, which has identified stable expression profiles/patterns [1,9,12,24-28]. Public repositories of gene expression data have appeared in the last decades. The Gene Expression Omnibus (GEO) [29], a well-known international public repository, stores and allows access to gene expression data generated by different high throughput technologies such as microarrays or next-generation sequencing. Exploiting and reusing the vast amount of data in these repositories has become a powerful tool for those searching for gene expression patterns across many diverse types of tissues and conditions.

A survey of forty studies of human adipose tissue (AT) published since 2001 noted that 70% of papers employed the *ACTB, GAPDH*, and *18S* HKGs as reference genes [14]. Related studies have supported the use of additional HKGs (i.e., *PPIA, HPRT, RPS18*, or *RPL19*) in human AT-based studies [16,30,31]. Importantly, these studies failed to include sex as a biological variable, suggesting that these HKGs may not be as suitable as anticipated. In short, there exists an important limitation in gene expression studies due to the lack of inclusion of the sex perspective. In response, this study determines the gene expression variability levels of six HKGs commonly used in human and mouse adipose tissue (AT) and genes included in various whole-transcriptome microarrays available at GEO that consider sex as a covariable. Further, we identify novel candidate reference genes that do not display sex bias in human AT. We extend this analysis to experimental analyses of mouse models deposited in the GEO. Our findings reveal that studies generally lack sex specificity or employ mainly male animals; furthermore, certain conventional HKGs fail the requisite of being constitutively expressed in both sexes. Also, we establish new putative sex unbiased HKGs (suHKGs) for gene expression analysis in male and female human AT, and putative orthologs for mouse AT. We present a general framework for reference gene selection that may be useful in gene expression studies and develop an open web tool to select adequate suHKGs according to customized experimental designs in AT.

## Methods

The bioinformatics analysis strategy was carried out using R 3.5.0 [32] and Python 3.0 and is summarized in **Fig. 1**.

**Fig. 1.**
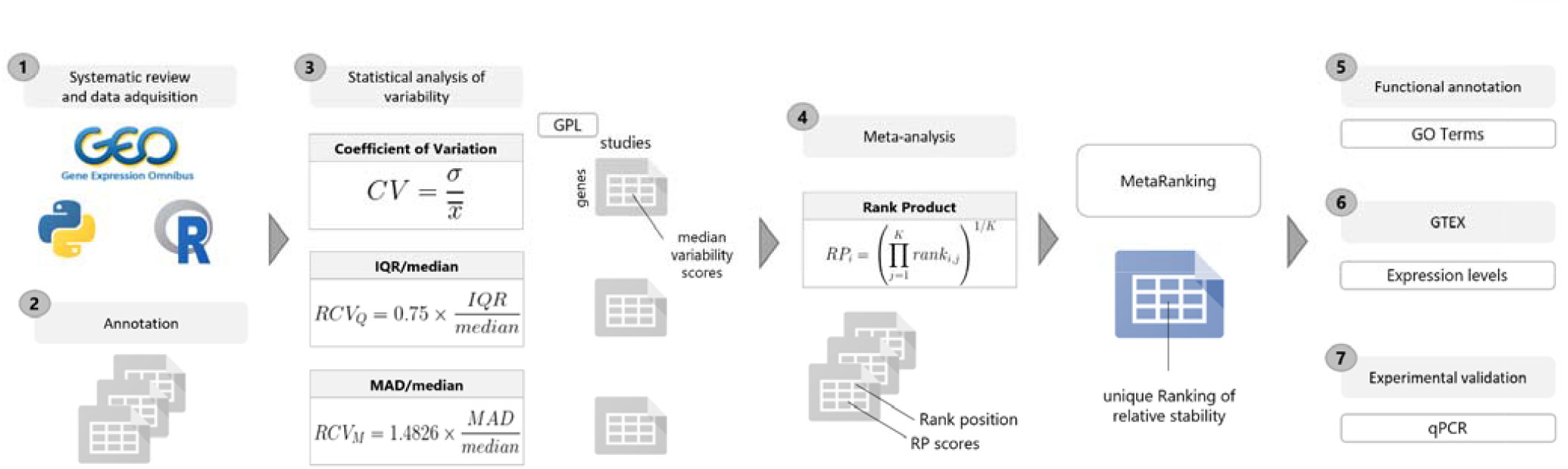
Data-analysis workflow. This study consisted of seven main block-steps: **1** The collection of public microarray information located at GEO (Gene Expression Omnibus) database with Python and R. **2** Raw data pre-processing and probe annotation. **3** Statistical data analysis with three different statistics to get the ene expression variability in adipose tissue samples of Hsa and Mmu, considering the biological sex as a variable. **4** Meta-analysis by Rank Product method. **5** Functional annotation with Gene Ontology (GO) terms. **6** GTEX-based gene expression filtering, to select potential reference genes suitable to compare both sexes in gene expression analyses. **7** Experimental validation by qPCR.

### Systematic Review and Data Collection

A comprehensive systematic review was conducted to identify all available transcriptomics studies with adipose tissue samples at GEO. The review considered the fields: sample source (adipose), type of study (expression profiling by array), and organism of interest (*Homo sapiens* or *Mus musculus*). The search was carried out during the first quarter of 2020, with the review period covering the years 2000-2019. From the returned records, the study GSE ID, the platform GPL ID, and the study type were extracted using the Python 3.0 library Beautiful Soup. The R package GEOmetadb [33] was then used to identify microarray platforms and samples from adipose tissue. The top 4 and 5 most used platforms in Hsa (**Table 1**) and Mmu (**Table 2**), respectively, were selected. Given the complex nature of some of the studies, those with information regarding the sex of samples were manually determined, and the keywords used to annotate them homogenized. Finally, studies not meeting the following predefined inclusion criteria were filtered out: i) include at least 10 adipose tissue samples, ii) use one of the selected microarray platforms to analyze gene expression data, iii) present data in a standardized way, and iv) not include duplicate sample records (as superseries).

**Table 1.** Processed data sets for selected studies of Hsa.

| **Platform** | **Description** | **Eligible Studies** | **Included Studies** | **Analyzed Samples** | **Identified Genes** |
| --- | --- | --- | --- | --- | --- |
| GPL570 | Affymetrix Human Genome U133 Plus 2.0 Array | 37 | 20 | 1058 | 22881 |
| GPL6244 | Affymetrix Human Gene 1.0 ST Array transcript (gene) version | 15 | 13 | 343 | 23307 |
| GPL10558 | Illumina HumanHT-12 V4.0 expression BeadChip | 14 | 7 | 498 | 31426 |
| GPL6947 | Illumina HumanHT-12 V3.0 expression BeadChip | 12 | 9 | 825 | 25159 |

The number of studies that used the platform (eligible studies) is shown for each selected platform, including the number of studies that met the exclusion criteria, the number of adipose tissue samples, and the maximum number of genes identified. For each selected platform, the number of studies that used the platform (eligible studies) are shown, including the number of studies that made the cut (refer exclusion criteria), the number of adipose tissue samples and the maximum number of genes that were able to be identified. A total of 49 studies and 2724 samples have been included in the statistical analysis.

**Table 2.** Processed data sets for selected Mmu studies.

| **Platform** | **Description** | **Eligible Studies** | **Included Studies** | **Analyzed Samples** | **Identified Genes** |
| --- | --- | --- | --- | --- | --- |
| GPL1261 | Affymetrix Mouse Genome 430 2.0 Array | 34 | 16 | 280 | 21495 |
| GPL6246 | Affymetrix Mouse Gene 1.0 ST Array transcript (gene) version | 24 | 6 | 133 | 24213 |
| GPL6887 | Illumina MouseWG-6 v2.0 expression BeadChip | 20 | 8 | 183 | 30886 |
| GPL6885 | Illumina MouseRef-8 v2.0 expression BeadChip | 15 | 8 | 375 | 18120 |
| GPL16570 | Affymetrix Mouse Gene 2.0 ST Array transcript (gene) version | 10 | 5 | 101 | 24647 |

The number of studies that used the platform (eligible studies) is shown for each selected platform, including the number of studies that met the exclusion criteria, the number of adipose tissue samples, and the maximum number of genes identified. 43 studies and 1072 samples have been included in the statistical analysis.

### Data Processing and Statistical Analyses

The normalized microarray expression data of the selected studies from GEO were downloaded using the GEOQuery R package. All the probe sets of each platform were converted to gene symbols, averaging expression values of multiple probe sets targeting the same gene to the median value.

Three statistical stability indicators were calculated for each gene in each study to determine the relative expression variability: the coefficient of variation (CV), the IQR/median, and the MAD/median. The CV, computed as the standard deviation divided by the mean, is used to compare variation between genes with expression levels at different orders of magnitude; however, extreme values can affect this value. Therefore, the interquartile range (IQR) divided by the median and the median absolute deviation (MAD) divided by the median (two statistics based on the median) were also considered. These measures provide more robustness in skewed distributions [34]. Both statistics were multiplied by a correction factor of 0.75 and 1.4826 to make them comparable to the CV in normal distributions. Lastly, the gene variability scores per platform were expressed as the median of all statistics from the studies analyzed with each platform. These median values were ranked by gene variability value for each platform, with lower ranks corresponding to higher stability levels.

The described analysis pipeline was performed on three different sample groups based on sex and species: female Hsa, male Hsa, and all Mmu samples. The analysis was not performed separately for male and female mice due to the lack of female Mmu samples.

### Meta-analysis

The gene variability ranks for each platform were integrated using the Rank Product (RP) method [35,36], a non-parametric statistic identifying the elements that systematically occupy higher positions in ranked lists. This approach combines gene ranks rather than variability scores to create platform independence. The RankProd package [37,38] was used to calculate the RP score for each gene (**Equation 1**, where *i* is the gene, *K* the number of platforms, and *rank*_*ij*_ the position of gene *i* in the ranking of platform *j*). Three final rankings were obtained (one for each sample group [Mmu, female, and Hsa male samples]) by sorting the genes in increasing order of RP.

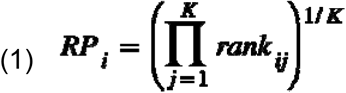

### Selection of Candidate HKGs

To encounter appropriate sex unbiased HKG (suHKG) candidates, male and female Hsa samples were randomly selected, and the Mmu group was discarded. Gene functional information was then incorporated to exclude genes involved in metabolic alterations. The AnnotationDbi and org.Hs.eg.db annotation packages converted Gene Symbol to Gene name. After removing pseudogenes and non-coding genes, the associated GO terms of the remaining genes were obtained using the GO.db annotation package. Related information from all three gene ontologies were included (Biological Process, Molecular Function, Cellular component). Genes related to physiopathological conditions were filtered out, and a unique ranking by sex was generated (the male and female *MetaRankings*), which averages the three statistical rankings (**Equation 2**).

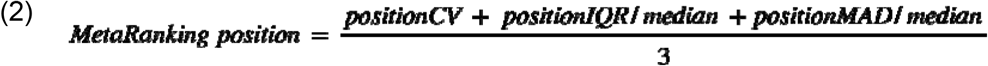

The difference in the ranking positions occupied in by males and females was also calculated to reveal sex-based stability differences at a gene level.

Selecting stable suHKG with high levels of expression, followed several steps-we first i) downloaded the “<u>GTEx_Analysis_2017-06-05_v8_RNASeQCv1.1.9_gene_median_tpm.gct.gz</u>” file from GTEx, ii) select the adipose tissue samples, iii) take the gene median transcript per million (TPM) value in visceral adipose tissue, iv) filter out from our sex-specific MetaRankings genes with median TPM < 20, v) select the genes in the top 10% positions of each MetaRanking, and vi) intersect the two top lis s to find stable and highly expressed genes common to both sexes.

## Experimental Validation

### Study Selection and Sample Processing

Subjects were recruited by the endocrinology and surgery departments at the University Hospital Joan XXIII (Tarragona, Spain) in accordance with the Helsinki declaration. Human visceral and subcutaneous AT samples were obtained during surgery from lean and obese male and female individuals. Total RNA was extracted from adipose tissue using the RNeasy lipid tissue midi kit (Qiagen Science). One microgram of RNA was reverse transcribed with random primers using the reverse transcription system (Applied Biosystems) [39].

Mouse AT was obtained from wild type and Irs2^-/-^ [40] (insulin resistance and type 2 diabetes model) C57BL/6 littermates. According to the criteria outlined in the “Guide for the Care and Use of Laboratory Animals,” all animals received humane care [22]. Total RNA was extracted from abdominal fat using a combined protocol including Trizol (Sigma) and RNeasy Mini Kit (Qiagen) with DNaseI Digestion. First-strand synthesis was performed using EcoDry Premix (Takara).

### Gene Expression Analysis

Quantitative gene expression analysis was performed on 50 ng c NA template. Real time-PCR was conducted in a LightCycler 480 Instrument IIR (Roche) using SYBR PreMix ExTaqTM (mi RNaseH Plus, Takara). Genes selected as potential HKG in human and mouse WAT were *18s, PPIA* and *RPL19*. Primers were designed in two consecutive exons, when possible, taking into consideration all reference sequences for mRNA in NCBI (https://www.ncbi.nlm.nih.gov/gene/; https://www.ncbi.nlm.nih.gov/nuccore/) [41] and aligned to search for common regions with Pairwise Sequence Alignment (https://www.ebi.ac.uk/Tools/psa/) **[42]**. Alternative transcript variants were analyzed by AceView (https://www.ncbi.nlm.nih.gov/IEB/Research/Acembly/index.html) [43] and primers (designed either by Primer3 or PrimerBlast) amplifying most represented sequence/s were chosen **(Additional file 1)**. All primers used in this study are noted in **Additional file 2**: **Table S1**. Crossing point (Cp) values were analyzed for stability between samples and relative quantification using 2^-ΔCt. Statistical analyses were performed with GraphPad Prism 8 (Graphpad Software V 8.0). The results are expressed as arithmetic mean ± the standard error of the mean (SEM). When two data sets were compared, a Student’s t-test was used. The differences observed were considered significant when: p-value <0.05 (*), p-value <0.01 (**) and p-value <0.001 (***).

### Web Tool

A freely available web tool, called metafun-HKG (https://bioinfo.cipf.es/metafun-HKG) was created during this study to allow users to review and share the large volume of generated data and results. The front-end was developed using the Bootstrap library. This easy-to-use resource is organized into four sections: **1**) a quick summary of the results obtained with the analysis pipeline in each phase. Then, for each of the studies, the detailed results of the **2**) exploratory analysis and **3**) variability assessment. Finally, all results are integrated and summarized in **4**) gene stability meta-analysis by sex and organism. The user can interact with the web tool through graphics and tables and search information for specific genes.

## Results

### Classic HKG selection

An extensive bibliographic review revealed that reference genes chosen for qRT-PCR-mediated analysis of gene expression in human AT or various types of adipocytes generally included the metabolic genes *GAPDH* [7,14-16,39,44], *HPRT* [7,16], *PPIA* [14,39,44], *UBC* [14,45] and ribosomal genes *18S* [7,14,16,39,46-48] and *RPL19* [49]. As these genes have been commonly used to analyze gene expression as reference genes in several experimental conditions (although the sex variable was generally not considered), we selected these six classic human AT HKG genes for evaluation when considering sex as a variable to assess their suitability as sex unbiased HKG (suHKGs). In the case of *18S*, we specifically selected *18S5* for our analysis.

### Systematic Review and Data Collection

We searched the GEO by defining the sample tissue, type of study, and organism of interest and obtained a total of 187 and 214 candidate studies for *Homo sapiens* (Hsa) and *Mus musculus* (Mmu), respectively. We selected the main microarray platforms for each species that contained the greatest number of studies; this provided 4 and 5 platforms for Hsa (**Table 1**) and Mmu (**Table 2**), respectively. We excluded 138 and 171 studies of Hsa and Mmu, respectively, as they failed to meet the inclusion criteria. Finally, we selected 49 Hsa studies and 43 Mmu studies for sex-based evaluations (**Fig. 2**), which involved 2,724 Hsa and 1,072 Mmu samples.

**Fig. 2.**
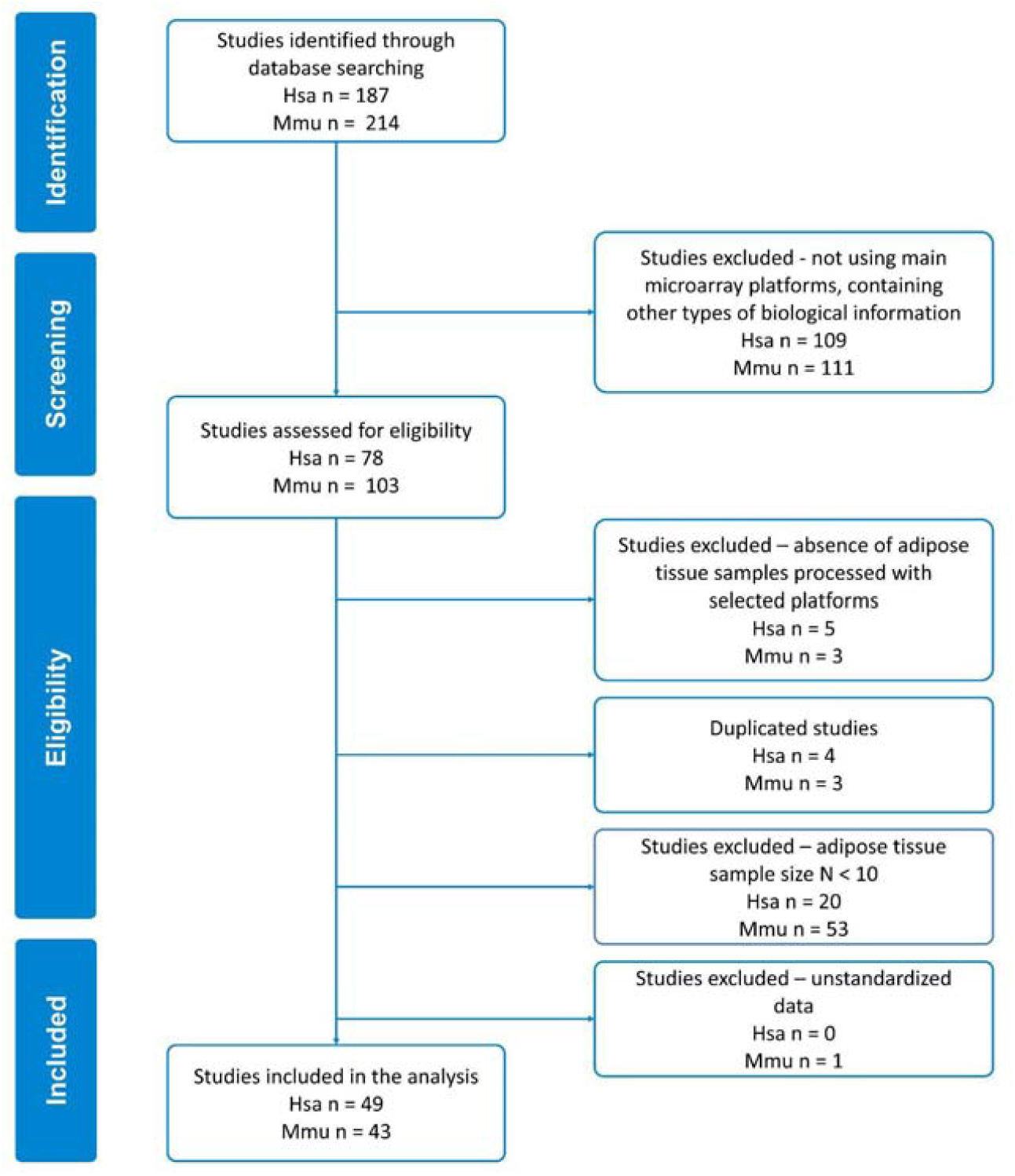
Flow diagram of the systematic review and selection of studies for meta-analysis according to PRISMA statement guidelines for database searches.

In Hsa, 24 (51%) of the 49 selected studies included sample information regarding sex. 10 studies covered both sexes in their analysis, while 11 included females exclusively, and 3 contained only male samples (**Fig. 3A**). In Mmu, 22 (51%) of the 43 selected studies informed about the sex of samples; only 1 study covered both sexes while 2 included exclusively female samples and 19 contained only male samples (**Fig. 3B**). Finally, we selected human samples with known sex information (681 male and 875 female samples, **Additional file 2: Table S2** and **Fig. S1**) and all mouse samples (1072 samples, 559 known to be male and 34 from female, **Additional file 2: Table S3** and **Fig. S2**) for analysis. Due to the low number of known female samples in mice, we excluded Mmu studies from this sex-based analysis.

**Fig. 3.**
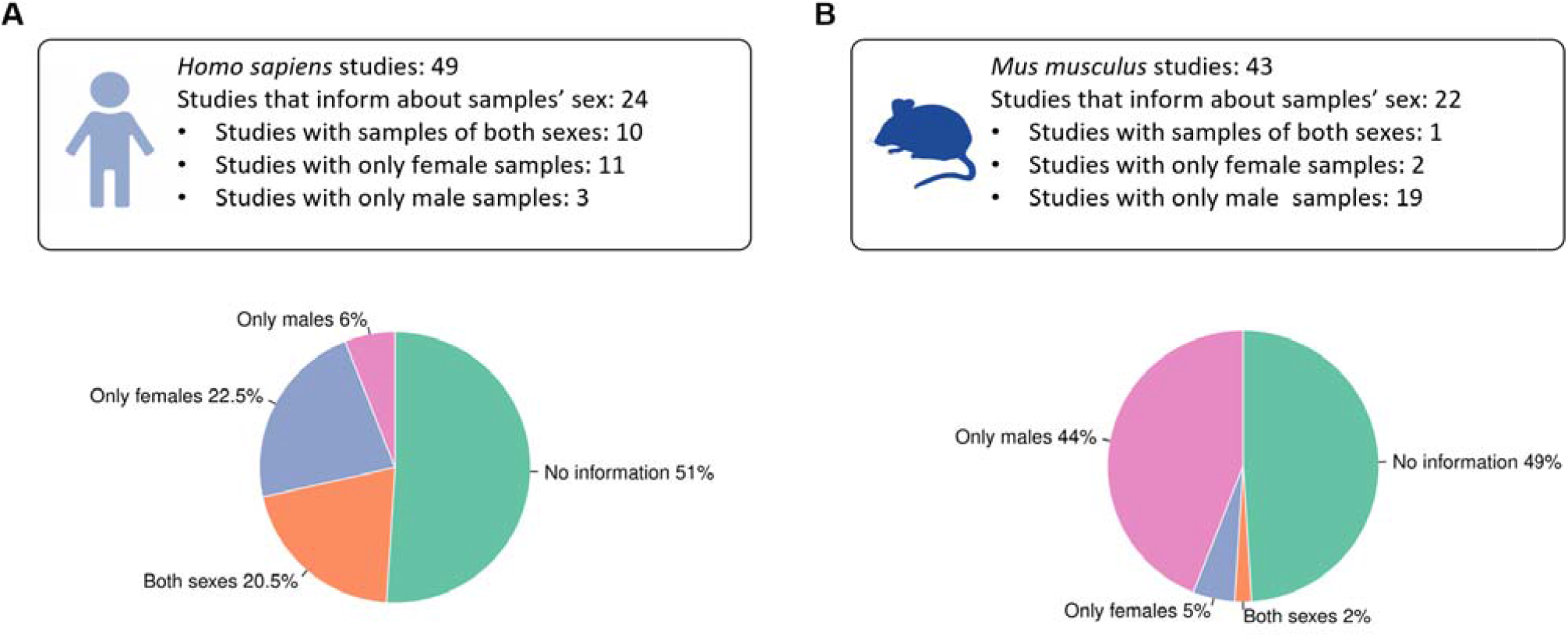
Summary of sex as a variable during the review of Hsa and Mmu studies. **A** Out of 49 Hsa studies, 49% specified the sex of samples, and 20.5% used samples from both sexes in the experimental procedure. **B** In Mmu, 51% of studies presented information regarding sex but focused mainly on male samples; almost no female samples were found in these studies. Only one study included samples from both sexes.

### Stability Data Meta-analysis

After downloading and annotating normalized expression data sets for the selected studies, we calculated three estimators of variability: the coefficient of variation (CV), the interquartile range divided by the median value (IQR/median), and the mean absolute deviation divided by the median value (MAD/median). **Additional file 2: Fig. S3, S4**, and **S5** summarize the levels of variability of the six selected HAT HKGs (*UBC, RPL19, RNA18S5, PPIA, HPRT1*, and *GAPDH*) for male and female Hsa and Mmu.

We conducted a meta-analysis based on the Rank Product (RP) method to integrate statistical results from different platforms; this approach combines gene ranks rather than variability scores (creating platform independence) and identifies the elements that systematically occupy higher positions in ranked lists (giving to each element in the ranking an RP score). We calculated the RP score of 41,975 and 47,203 Hsa and Mmu genes, respectively, and then sorted all genes-in this ranking, lower positions indicate higher expression stability. *18S* displayed significant variability in Hsa in both males and females; however, this gene represented the second most stable selected HKG in Mmu. **Fig. 4** depicts the positions occupied by the six selected HAT HKGs in Mmu, Hsa males, and Hsa females. Surprisingly, HKG stability in humans differed between female and male samples, with females displaying greater instability. Accessing the Metafun-HKG webtool provides the whole rankings with the positions and RP scores of all evaluated genes in each experimental condition.

**Fig. 4.**
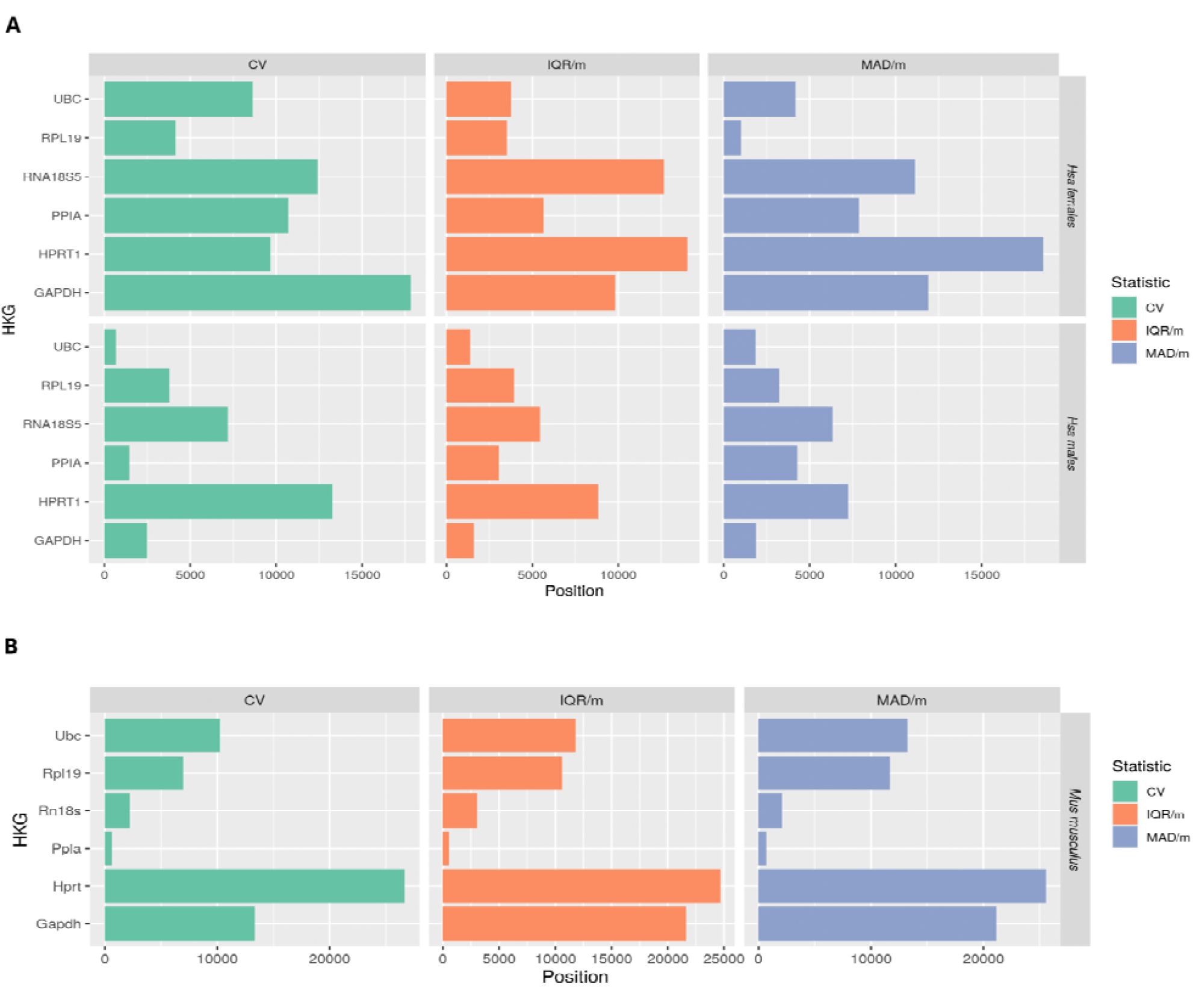
A Ranking of stability levels for classic HKGs evaluated in Hsa females (upper) and males (lower). The position in the ranking for each selected gene is described on the X-axis. This ranking was generated by taking the mean of the obtained RP values for the three statistical approaches (CV, IQR/median, and MAD/median) after filtering non-coding genes. Ranking based on 18973 genes. **B** Ranking stability levels for classic HKGs evaluated in Mmu. This ranking was generated by taking the mean of the obtained RP values for the three statistical approaches (CV, IQR/median, and MAD/median). Ranking based on 47203 genes.

In order to decipher differences in gene expression stability between male and female AT samples, we conducted a deconvolution analysis. Overall, we did not find consistent differences between the sexes in cellular composition across datasets (**Additional file 3**).

To select sex-unbiased, highly-expressed, and stable human human AT HKG candidates, we combined the scores of the three statistical approaches in a unique list of positions for each experimental condition (metaRanking) and filtered out genes with low expression (TPM < 20) in the GTEx database. These steps provided a list of 5,315 genes. We next intersected the top 10% (532) most stable genes in the Hsa male and Hsa female metaRankings separately, which resulted in a list of 195 candidate suHKGs (http://bioinfo.cipf.es/metafun-HKG/). This analysis revealed relative stability and expression values high enough for detection by different gene expression analysis technologies in total Hsa samples (**Table 3, Fig. 5**). From this list, we selected human AT HKGs that included the classical HKGs *PPIA, UBC, RPL19*, and *RPS18* and the additional novel candidate suHKGs *RPS8* and *UBB*. We also detected stable, highly-expressed genes in one sex but not in the other (such genes included *ANXA2, DDX39B*, and *PLIN4* in males and *DNASE2, NDUFB11*, and *RARA* in females (**Additional file 2: Table S4, Fig. 5**), which may be used as sex-specific reference genes. We failed to find the expression of the *18S* gene in GTEx, although we searched for different aliases *(RNA18S5, RNA18S1, RNA18SN1, RNA18SN5, RN18S1)*.

**Table 3.** Candidate suHKGs for gene expression analysis.

| Gene | Relative stability in male | Relative stability in female | Expression level (TPM) | Expression level (TPM) in female | Expression level (TPM) in male |
| --- | --- | --- | --- | --- | --- |
| *PPIA* | 873 | 1,589 | 234.597 | 236.1 | 233.6 |
| *RPL19* | 1,129.67 | 137.33 | 1,707.61 | 1707 | 1708 |
| *RPS8* | 194.67 | 178.33 | 952.191 | 944.5 | 957.9 |
| *RPS18* | 119.33 | 296.33 | 3,173.82 | 3180 | 3168 |
| *UBB* | 228 | 79 | 252.293 | 249.8 | 254.1 |
| *UBC* | 267.33 | 706.33 | 432.547 | 396.9 | 447.7 |

Selection of housekeeping candidate genes proposed to be used as a reference to compare both sexes in gene expression analysis. *PPIA* and *RPL19* have been experimentally validated, while *RPS8*, *RPS18*, *UBB*, and *UBC* are computationally suggested. These genes are proposed based on their sex-specific values of relative expression stability obtained from the final MetaRanking positions. Expression levels have been extracted from GTEx (given in TPM), which are high enough for detection by different technologies.

**Fig. 5.**
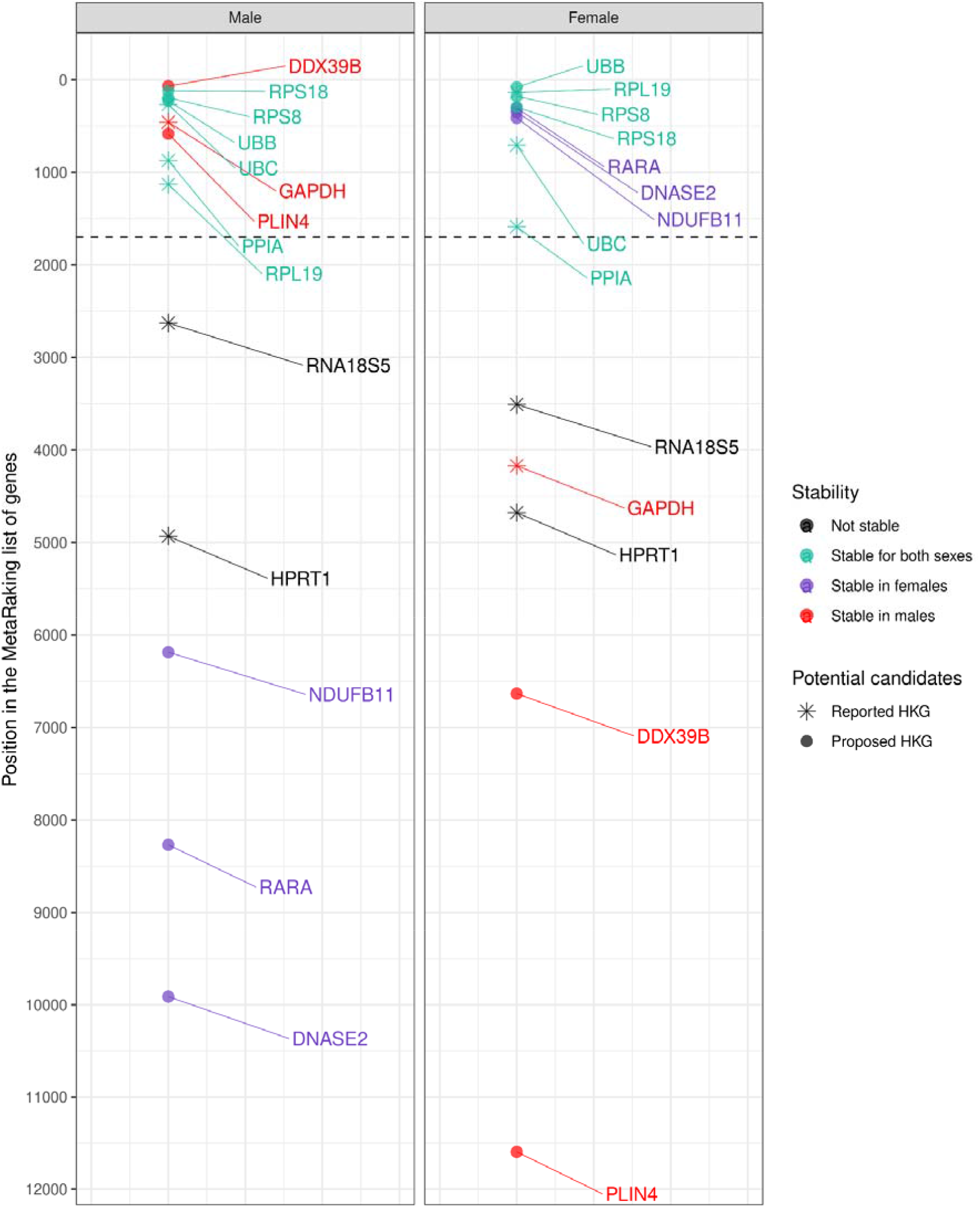
MetaRanking of HKG stability levels for Hsa females and males. Dot shape indicates classical HKG (star) or new potential HKGs (circle). The color indicates if a gene is stable for both sexes (green), only in females (violet), only in males (red), or unstable (black). Dashed line indicates the limit position of the top 10% most stable genes with an expression of at least 20 TPM.

### Experimental Validation

We selected *PPIA, RPL19*, and *18S* for experimental validation according to our computational assessment of variability. We analyzed human AT mRNA from lean and obese male and female individuals by qPCR to validate the previous computational metadata analysis (**Table 3; Fig. 6**). Raw crossing point (Cp) value coefficient variation (CV) analysis revealed similar Cp values between male and female samples, with low CV values for *PPIA* and *RPL19* (**Fig. 6A**); however, *18S* exhibited significant differences in Cp values between male and female samples, which displayed high CV values (**Fig. 6A**). Further, gene expression analysis of multiple experimental targets revealed differing patterns when using *PPIA* or *RPL19* compared to *18S* as a HKG (**Fig. 6B**). We analyzed several genes involved in physiological and metabolic adipose tissue functions (e.g., *IRS1, LEPR*, and *PPAR*_Y_) in male and female human AT samples under two different physiological conditions using potential suHKG candidates. Results obtained provided evidence for the suitability of *RPL19* and *PPIA* as suHKGs and disqualified *18S* as a HKG when considering sex as a variable (**Fig. 6B**). Overall, the experimental procedures validate the computational metadata analysis, discarding *18S* and selecting *PPIA* and *RPL19* as suHKG for HAT analysis.

**Fig. 6.**
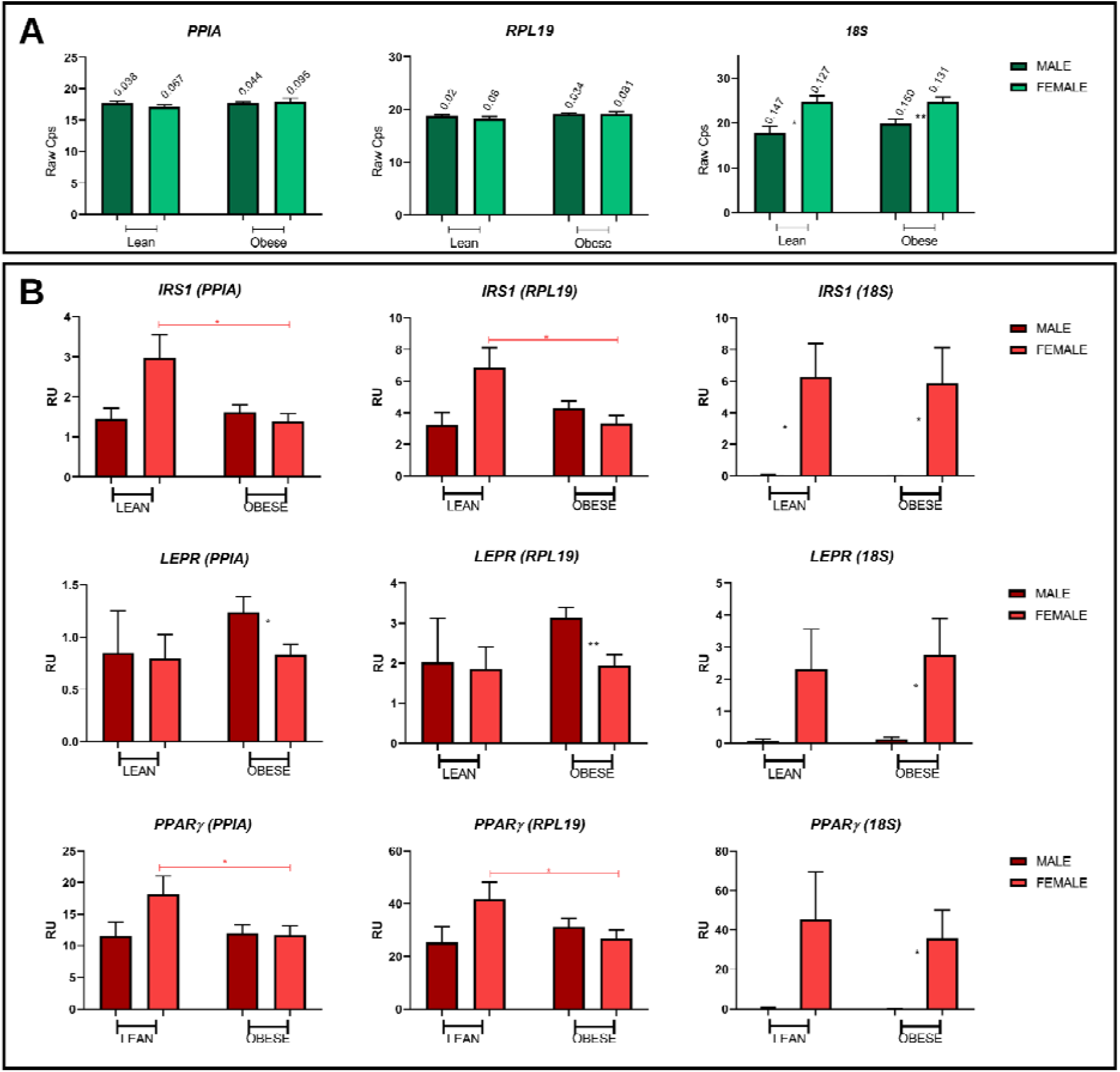
Gene expression analysis in HAT from male and female samples using different HKGs. **A** Coefficient of variation (CV) in the Cp values of each candidate gene calculated in male and female for lean and obese samples. **B** *IRS1, LEPR*, and *PPAR* expression analysis using *PPIA, RPL19*, and *18S* as reference genes. Male Lean n=3; Female Lean n=7; Male Obese n=10; Female Obese n=10. Student’s t-test applied for significance-(*) p-value<0.05, and (**) p-value<0.01.

To circumnavigate the lack of sex-based Mmu data to compute a Mmu metaRanking, we experimentally evaluated mouse orthologs (*Ppia, Rpl19*, and *18s*) of validated human suHKGs, in wt and in an insulin resistance, Irs2^-/-^ ko model in male and females. Relative gene expression analysis demonstrated that the internal control affected the relative expression of different experimental targets in different experimental mouse models. *18s* used as HKG alters relative gene expression of *InsR, Lepr*, and *Phb* in males and females mouse AT samples, while *Ppia* and *Rpl19* succeed as suHKG in mouse AT samples. (**Additional file 2: Fig. S6**). These results confirm that mouse homologs of suHKG candidates can be used in mouse-based gene expression studies.

### Metafun-HKG Web Tool

We created the open platform web tool Metafun-HKG (https://bioinfo.cipf.es/metafun-HKG) to allow easy access to any information related to this study. This resource contains information related to the study samples, systematic revision, gene variability scores, and stability rankings. The stability indicators for each gene evaluated by platform, species, and sex can be freely explored by users to identify profiles of interest.

## Discussion

### Assessment of suHKG Candidates

The two main objectives of this work were i) evaluating the suitability of a group of six classic HKGs acting as human AT suHKGs and ii) identifying genes with a stable, high expression profile that represent new Human AT suHKG candidates. Our novel strategy has reviewed the role of HKGs by considering sex, species, and platform as variables in evaluated studies.

We performed our analysis on three different sample groups based on sex and species: female Hsa, male Hsa, and all Mmu samples. We did not analyze Mmu female and male samples separately due to the lack of reported female Mmu samples in the selected studies. HKGs displayed platform-dependent variability under all conditions, given that each microarray platform has its probe design and technical protocol. Previous studies on technology dependence concluded that this factor has less determining power than the differences in transcript expression levels caused by varying cell conditions [24].

Results exhibit considerable differences in gene stability, including stability differences in the six classical selected HKGs between Hsa female and male samples showing higher instability in females in general term, although deconvolution assays show no major differences in cell composition between males and females. *PPIA, UBC*, and *RPL19* displayed high stability levels for samples from both sexes, while *HPRT1* and *18S* exhibited low stability levels in both sexes. Interestingly, *GAPDH* displayed high stability in male samples and low stability in female samples. In apparent contradiction, *18s* presents high stability levels in Mmu, but this may be explained by the overwhelming presence of male samples in this group and the fact that this gene suffers a significant sex bias in mouse (**Additional file 2: Fig. S6**). The common absence of female samples in studies (as further evidenced by our systematic review) could explain the systematic reports of *18s* as a stable HKG.

We propose a list of 195 suHKG candidates suitable for use as internal controls in HAT-based gene expression studies including male and female samples; these genes exhibit high expression (TPM > 20) and stability levels and a minimal influence of sex on expression patterns. As we could not reproduce the pipeline followed with human samples in mouse studies due to the lack of female mouse samples, we suggest the orthologs of proposed human suHKGs as mouse suHKGs.

We validated a selection of suHKG candidates experimentally to assess the robustness of our computational findings; overall, our gene expression analysis validated the *in silico* results (**Table 3**). *PPIA*, a widely used human AT HKG, and *RPL19*, used as a HKG in several cell types [30,31,50] and occasionally in human AT studies [49], have been validated as human AT suHKGs; however, experimental validation shows that *18S*, which is widely used as human AT HKG [7,14,16,39,46-48], displays significant levels of variability in both male and female samples and sex-specific expression patterns (**Fig. 6**). These results agree with the findings of other recently published studies [51] and correlate with those found in mouse adipose tissue. The use of *18s* as a HKG induces apparent differences in the relative expression levels of several genes in males and females and wild type and Irs2^-/-^ samples (**Additional file 2: Fig. S6**); instead, we suggest *Rpl19* and *Ppia* as more optimal suHKGs in mouse adipose tissue analysis.

We identified several additional genes human AT suHKGs from the computational analysis, including *RPS18, RPS8*, and *UBB* (**Table 3**), that present characteristics such as appropriate stable and high expression levels. We also suggest the mouse orthologs of these human suHKGs as mouse suHKGs. To this end, we designed a web tool to customize the best suHKG for human or mouse adipose tissue experimental design.

### Strengths and Limitations

Massive data analysis of gene expression represents a pivotal tool for understanding different biological scenarios, which may eventually help elucidate mechanisms affecting basic and biomedical research. Data analyses must be assessed in the laboratory by studying relative gene expression normalized to an adequately chosen HKG. Selection of an ideal HKG remains a challenging process, although this choice will help to ensure an accurate result and must consider all experimental conditions and biological variables. Incorporating sex-based analyses into research will improve reproducibility and experimental efficiency by influencing the outcome of experiments and must be accounted for as a critical biological variable. Sex must be considered to monitor sex-based differences and similarities for all diseases and biological processes that affect both sexes, which may help reduce bias, enable social equality in scientific outcomes, and encourage new opportunities for discovery and innovation, as evidenced by several studies analyzing this issue [20,22,52-55].

Numerous lines of evidence suggest that the current status quo does not address fundamental issues of sex-based differences evident in gene expression. Up to date, many classic HKGs remain unevaluated when including sex as a biological variable; these include those commonly used in human AT studies (e.g., *ACTB, GAPDH*, and *18S*) and additional HKGs such as *PPIA, HPRT, RPS18*, or *RPL19*. Using a HKG to normalize samples without assessing their behavior under the specific experimental conditions used in each study (including sex), may lead to a biased outcome. HKGs may remain stable in one sex but not in the other, as in the case of *DDX39B* and *PLIN4* (stable in males) or *NDUFB11* and *RARA* (stable in females), or may have stable yet distinct expression levels in both sexes, such as for *18s* in mouse. Ignoring sex and choosing a non-optimal HKG may introduce confounding variables and the inability to assess whether differences in the data derived from the experimental design or the normalization process. This source of variability in the data would reduce statistical power, thereby making it more difficult to find significant results. In this study, we analyzed the role of six conventional HAT HKG considering sex as a variable for the first time.

Many published studies do not include a sex-based perspective by omitting animal sex from reporting of the animals or performing studies with animals of only one sex (typically males). Our systematic review found that 51% of Hsa studies and 49% of Mmu studies failed to include information regarding the sex of samples, with just 19% of Hsa and a striking 2% of Mmu studies including samples from both sexes. Of note, Mmu studies including only female samples represented just 5% of the total. The small number of Mmu studies, including female sample information, represented a significant limitation of the study and prevented the creation of a Mmu meta-ranking to select highly-expressed stable Mmu suHKG candidates as for Hsa. We evaluated the Mmu orthologs of the selected Hsa suHKG candidates experimentally to overcome this limitation, which confirmed their suitability as Mmu suHKGs.

Despite the widespread use of *18S* RNA as a HKG, its annotation represents another limiting factor of this study; we failed to encounter this gene in the GTEx platform under any proposed alias from GeneCards. We also noted that identifiers for this gene are unstable or not included in reference assemblies. In addition, the DNA sequence of the *RNA18SN5* gene (accession number NR_003286.4) has 99-100% identity with other ribosomal RNAs such as *RNA18SN1, RNA18SN2, RNA18SN3, RNA18SN4*, and *RNA18SP3* (accession numbers NR_145820.1, NR_146146.1, NR_146152.1, NR_146119.1, NG_054871.1, respectively). Furthermore, *18S* rRNA has different copy numbers among individuals and varies with age [56]. Considering all these factors, and integrating experimental data assessing differential expression levels according to sex, makes the *18S* gene less suitable as a HAT suHKG than other suHKGs proposed in this study.

Other limitations of the study included the filtering and pre-processing of biological information located in the GEO to identify the published studies with transcriptomic data of adipose tissue, and the classification of the samples depending on the sex. A primary limiting factor involved the absence of standardized vocabulary to tag sex in sample records of the studies. Even though the gene expression data in GEO is presented as a standardized expression matrix, the metadata (including sample source, tissue type, or sample sex) is reported through free-text fields written by the researcher submitting the study. The absence of standardized vocabulary and structured information constrains data mining power on large-scale data, and improvements in this regard could aid the processing of data in public repositories [57].

For the first time, this study presents a computational strategy that includes a massive data analysis capable to assess the sex bias in expression levels of classical and novel HKGs, over a large volume of studies and samples. This strategy revealed that an accurate experimental design for adipose tissue requires the adequate selection of a suHKG, such as *PPIA, RPL19*, or new options, such as *RPS18* or *UBB*. In that context, we could finally avoid the common practice of pooling males and females or even discard the only male-presence effect. This study presents the relative expression stability of six commonly used HKGs and the variability levels of other genes covered by the analyzed microarray platforms. This strategy is aligned with the FAIR principles [58] (Findability, Accessibility, Interoperability, and Reusability) to ensure the further utility and reproducibility of the generated information.

Although limited to adipose tissue, our findings suggest that the sex bias in commonly used HKGs could appear in other tissues, thereby affecting the normalization process of gene expression analysis of any kind. Incorrect normalization may significantly alter gene expression data, as shown in the case of 18S, and lead to erroneous conclusions. This study highlights the importance of considering sex as a variable in biomedical studies and provides evidence that thorough analyses of HKGs as internal controls in all tissues should be promptly addressed.

### Perspectives and Significance

Our results focus on the importance of taking into consideration sex as a biological variable when choosing the best HKG as reference in HAT gene expression analysis. Our novel computational strategy includes massive data analysis capable to assess the sex bias in expression levels of classical and novel HKGs to select sex-unbiased HKG. Conventionally reported HKG genes include several metabolic and ribosomal genes such as GAPDH, HPRT, PPIA, UBC, 18S and RPL19. However, our novel computational strategy based on meta-analysis techniques has proven that certain classical HKGs, like one of the most extended, 18S, may fail to function adequately as the reference gene as it differentially expressed in males and females, while others like PPIA and RPL19, succeeded as reference genes. Further, following selection criteria, several markers, like RPS8 and UBB are also proposed and an open web resource (https://bioinfo.cipf.es/metafun-HKG) offered for customized experimental design.

All these results provide new useful insight in evaluating gene expression analysis in human adipose tissue under several experimental conditions and with biomedical purposes. Using an incorrect HKG may lead to inappropriate results interpretation and applications, while using a suHKG will always provide a better experimental approach, either when taking into consideration male and females as separate groups, either included in the same experimental group but properly analyzed. This study highlights the importance of considering sex as a variable in gene expression analyses in human AT and provides evidence for future extensive tissues suHKG selection to be hopefully, promptly addressed.

## Supplementary information

**Additional file 1**. HKG primers design process.

**Additional file 2. Table S1**. List of primers used for the experimental validation. **Table S2**. Human sample information. **Table S3**. Mouse sample information. **Table S4**. Selection of candidate sex-specific HKGs in gene expression analysis. **Fig. S1**. Summary of the number of female and male samples found in each Hsa study. **Fig. S2**. Summary of the number of female and male samples found in each Hsa study. **Fig. S3**. Variability levels for classic HKGs evaluated in Hsa females. **Fig. S4**. Variability levels for classic HKGs evaluated in Hsa males. **Fig. S5**. Variability levels for classic HKGs evaluated in all Mmu samples. **Fig. S6**. Candidate HKG analysis in mouse adipose tissue, using wt and irs2^-/-^ KO male and female samples.

**Additional file 3**. Methods and results of deconvolution analysis.

## Acknowledgments

The authors thank the Principe Felipe Research Center (CIPF) for providing access to the computer cluster. Part of the equipment employed in this work has been funded by Generalitat Valenciana and co-financed with ERDF funds (OP ERDF of Comunitat Valenciana 2014-2020). The authors also thank Stuart P. Atkinson for reviewing the manuscript.

## Data Availability

All datasets used in this work are available in the Gene Expression Omnibus public repository.

## Author contributions statement

Conceptualization, F.G.G., and A.G.; methodology, M.G., R.G.R., M.R.H., A.G., and F.G.G.; software, M.G., and M.R.H.; formal analysis, M.G., B.G.C., G.A.B., H.G.M., and R.G.R.; investigation, M.G., R.G.R., M.R.H., A.G., and F.G.G.; data curation, M.G., and R.G.R.; experiment conduction: A.G. and S.F.V.; writing—original draft preparation, M.G., R.G.R., M.R.H., D.B., S.F.V., A.G., and F.G.G.; writing—review and editing, M.G., R.G.R., M.R.H., A.G., and F.G.G.; visualization, M.G., R.G.R., M.R.H.; supervision, A.G., M.R.H., and F.G.G.; funding acquisition, M.G., F.G.G., and D.B.; project administration, F.G.G., M.R.H., and A.G. All authors have read and agreed to the published version of the manuscript.

## Declarations

### Consent for publication

Not applicable.

### Ethics approval and consent to participate

The study was conducted according to the guidelines of the Declaration of Helsinki, and approved by the Ethics Committee of IISPV (Tarragona, Spain). Informed consent was obtained from all subjects involved in the study. Subjects were recruited by the Endocrinology and Surgery departments at the University Hospital Joan XXIII. The samples were stored in a tissue biobank registered at the National Register of Biobanks (registration number #C.0003609). Committee Reference Code: CEIm 34p/2016.

## References

1. Chang CW, Cheng WC, Chen CR, et al. Identification of Human Housekeeping Genes and Tissue-Selective Genes by Microarray Meta-Analysis. PLOS ONE. 2011;6(7):e22859. doi:10.1371/journal.pone.0022859

2. Caracausi M, Piovesan A, Antonaros F, Strippoli P, Vitale L, Pelleri MC. Systematic identification of human housekeeping genes possibly useful as references in gene expression studies. Mol Med Rep. 2017;16(3):2397–2410. doi:10.3892/mmr.2017.6944

3. Eisenberg E, Levanon EY. Human housekeeping genes, revisited. Trends Genet. 2013;29(10):569–574. doi:10.1016/j.tig.2013.05.010

4. Butte AJ, Dzau VJ, Glueck SB. Further defining housekeeping, or “maintenance,” genes Focus on “A compendium of gene expression in normal human tissues.” Physiol Genomics. 2001;7(2):95–96. doi:10.1152/physiolgenomics.2001.7.2.95

5. Dheda K, Huggett JF, Bustin SA, Johnson MA, Rook G, Zumla A. Validation of housekeeping genes for normalizing RNA expression in real-time PCR. BioTechniques. 2004;37(1):112–119. doi:10.2144/04371RR03

6. Bustin SA, Benes V, Garson J, et al. The need for transparency and good practices in the qPCR literature. Nat Methods. 2013;10(11):1063–1067. doi:10.1038/nmeth.2697

7. Chechi K, Gelinas Y, Mathieu P, Deshaies Y, Richard D. Validation of Reference Genes for the Relative Quantification of Gene Expression in Human Epicardial Adipose Tissue. PLOS ONE. 2012;7(4):e32265. doi:10.1371/journal.pone.0032265

8. Stürzenbaum SR, Kille P. Control genes in quantitative molecular biological techniques: the variability of invariance. Comp Biochem Physiol B Biochem Mol Biol. 2001;130(3):281–289. doi:10.1016/S1096-4959(01)00440-7

9. She X, Rohl CA, Castle JC, Kulkarni AV, Johnson JM, Chen R. Definition, conservation and epigenetics of housekeeping and tissue-enriched genes. BMC Genomics. 2009;10(1):269. doi:10.1186/1471-2164-10-269

10. Zhu J, He F, Song S, Wang J, Yu J. How many human genes can be defined as housekeeping with current expression data? BMC Genomics. 2008;9(1):172. doi:10.1186/1471-2164-9-172

11. Suzuki T, Higgins PJ, Crawford DR. Control Selection for RNA Quantitation. BioTechniques. 2000;29(2):332–337. doi:10.2144/00292rv02

12. Hsiao LL, Dangond F, Yoshida T, et al. A compendium of gene expression in normal human tissues. Physiol Genomics. 2001;7(2):97–104. doi:10.1152/physiolgenomics.00040.2001

13. Jonge HJM de, Fehrmann RSN, Bont ESJM de, et al. Evidence Based Selection of Housekeeping Genes. PLOS ONE. 2007;2(9):e898. doi:10.1371/journal.pone.0000898

14. Gabrielsson BG, Olofsson LE, Sjögren A, et al. Evaluation of reference genes for studies of gene expression in human adipose tissue. Obes Res. 2005;13(4):649–652. doi:10.1038/oby.2005.72

15. Heo JS, Choi Y, Kim HS, Kim HO. Comparison of molecular profiles of human mesenchymal stem cells derived from bone marrow, umbilical cord blood, placenta and adipose tissue. Int J Mol Med. 2016;37(1):115–125. doi:10.3892/ijmm.2015.2413

16. White JM, Piron MJ, Rangaraj VR, Hanlon EC, Cohen RN, Brady MJ. Reference Gene Optimization for Circadian Gene Expression Analysis in Human Adipose Tissue. J Biol Rhythms. 2020;35(1):84–97. doi:10.1177/0748730419883043

17. Karp NA, Mason J, Beaudet AL, et al. Prevalence of sexual dimorphism in mammalian phenotypic traits. Nat Commun. 2017;8(1):15475. doi:10.1038/ncomms15475

18. Oliva M, Muñoz-Aguirre M, Kim-Hellmuth S, et al. The impact of sex on gene expression across human tissues. Science. 2020;369(6509). doi:10.1126/science.aba3066

19. Yoon DY, Mansukhani NA, Stubbs VC, Helenowski IB, Woodruff TK, Kibbe MR. Sex bias exists in basic science and translational surgical research. Surgery. 2014;156(3):508–516. doi:10.1016/j.surg.2014.07.001

20. Tannenbaum C, Ellis RP, Eyssel F, Zou J, Schiebinger L. Sex and gender analysis improves science and engineering. Nature. 2019;575(7781):137–146. doi:10.1038/s41586-019-1657-6

21. Woitowich NC, Beery A, Woodruff T. A 10-year follow-up study of sex inclusion in the biological sciences. Sugimoto C, Rodgers P, Shansky R, Schiebinger L, eds. eLife. 2020;9:e56344. doi:10.7554/eLife.56344

22. McCullough LD, de Vries GJ, Miller VM, Becker JB, Sandberg K, McCarthy MM. NIH initiative to balance sex of animals in preclinical studies: generative questions to guide policy, implementation, and metrics. Biol Sex Differ. 2014;5:15. doi:10.1186/s13293-014-0015-5

23. Accounting for sex and gender makes for better science. Nature. 2020;588(7837):196–196. doi:10.1038/d41586-020-03459-y

24. Lee PD, Sladek R, Greenwood CMT, Hudson TJ. Control Genes and Variability: Absence of Ubiquitous Reference Transcripts in Diverse Mammalian Expression Studies. Genome Res. 2002;12(2):292–297. doi:10.1101/gr.217802

25. Lee SR, Jo MJ, Lee JE, Koh SS, Kim SY. Identification of Novel Universal Housekeeping Genes by Statistical Analysis of Microarray Data. BMB Rep. 2007;40(2):226–231. doi:10.5483/BMBRep.2007.40.2.226

26. Popovici V, Goldstein DR, Antonov J, Jaggi R, Delorenzi M, Wirapati P. Selecting control genes for RT-QPCR using public microarray data. BMC Bioinformatics. 2009;10(1):42. doi:10.1186/1471-2105-10-42

27. Pilbrow AP, Ellmers LJ, Black MA, et al. Genomic selection of reference genes for real-time PCR in human myocardium. BMC Med Genomics. 2008;1(1):64. doi:10.1186/1755-8794-1-64

28. Zhang Y, Li D, Sun B. Do Housekeeping Genes Exist? PLoS ONE. 2015;10(5). doi:10.1371/journal.pone.0123691

29. Home - GEO - NCBI. Accessed December 1, 2021. https://www.ncbi.nlm.nih.gov/geo/

30. Galan A, Diaz-Gimeno P, Poo ME, et al. Defining the Genomic Signature of Totipotency and Pluripotency during Early Human Development. PLOS ONE. 2013;8(4):e62135. doi:10.1371/journal.pone.0062135

31. Galán A, Simón C. Monitoring Stemness in Long-Term hESC Cultures by Real-Time PCR. In: Turksen K, ed. Human Embryonic Stem Cell Protocols. Methods in Molecular Biology. Humana Press; 2010:135–150. doi:10.1007/978-1-60761-369-5_8

32. Team R. Core. R: A language and environment for statistical computing. 2013.:16.

33. Zhu Y, Davis S, Stephens R, Meltzer PS, Chen Y. GEOmetadb: powerful alternative search engine for the Gene Expression Omnibus. Bioinformatics. 2008;24(23):2798–2800. doi:10.1093/bioinformatics/btn520

34. Arachchige C, Prendergast L, Staudte R. Robust analogs to the coefficient of variation. J Appl Stat. Published online August 20, 2020:1–23. doi:10.1080/02664763.2020.1808599

35. Breitling R, Armengaud P, Amtmann A, Herzyk P. Rank products: a simple, yet powerful, new method to detect differentially regulated genes in replicated microarray experiments. FEBS Lett. 2004;573(1-3):83-92. doi:https://doi.org/10.1016/j.febslet.2004.07.055

36. Mitchell L. A parallel implementation of the Rank Product method for R. :12.

37. Hong F, Breitling R, McEntee CW, Wittner BS, Nemhauser JL, Chory J. RankProd: a bioconductor package for detecting differentially expressed genes in meta-analysis. Bioinformatics. 2006;22(22):2825–2827. doi:10.1093/bioinformatics/btl476

38. Del Carratore F, Jankevics A, Eisinga R, Heskes T, Hong F, Breitling R. RankProd 2.0: a refactored bioconductor package for detecting differentially expressed features in molecular profiling datasets. Bioinformatics. 2017;33(17):2774–2775. doi:10.1093/bioinformatics/btx292

39. Ceperuelo-Mallafré V, Duran X, Pachón G, et al. Disruption of GIP/GIPR axis in human adipose tissue is linked to obesity and insulin resistance. J Clin Endocrinol Metab. 2014;99(5):E908–919. doi:10.1210/jc.2013-3350

40. Withers DJ, Gutierrez JS, Towery H, et al. Disruption of IRS-2 causes type 2 diabetes in mice. Nature. 1998;391(6670):900–904. doi:10.1038/36116

41. Sayers EW, Bolton EE, Brister JR, et al. Database resources of the national center for biotechnology information. Nucleic Acids Res. 2022;50(D1):D20–D26. doi:10.1093/nar/gkab1112

42. Needleman SB, Wunsch CD. A general method applicable to the search for similarities in the amino acid sequence of two proteins. J Mol Biol. 1970;48(3):443–453. doi:10.1016/0022-2836(70)90057-4

43. Thierry-Mieg D, Thierry-Mieg J. AceView: a comprehensive cDNA-supported gene and transcripts annotation. Genome Biology. 2006;7(1):S12. doi:10.1186/gb-2006-7-s1-s12

44. Amisten S. Quantification of the mRNA expression of G protein-coupled receptors in human adipose tissue. Methods Cell Biol. 2016;132:73–105. doi:10.1016/bs.mcb.2015.10.004

45. Su X, Yao X, Sun Z, Han Q, Zhao RC. Optimization of Reference Genes for Normalization of Reverse Transcription Quantitative Real-Time Polymerase Chain Reaction Results in Senescence Study of Mesenchymal Stem Cells. Stem Cells Dev. 2016;25(18):1355–1365. doi:10.1089/scd.2016.0031

46. Ebrahimi R, Bahiraee A, Alipour NJ, Toolabi K, Emamgholipour S. Evaluation of the Housekeeping Genes; β-Actin, Glyceraldehyde-3-Phosphate-Dehydrogenase, and 18S rRNA for Normalization in Real-Time Polymerase Chain Reaction Analysis of Gene Expression in Human Adipose Tissue. Arch Med Lab Sci. 2018;4(3). doi:10.22037/amls.v4i3.26269

47. Gómez-Abellán P, Díez-Noguera A, Madrid JA, Luján JA, Ordovás JM, Garaulet M. Glucocorticoids Affect 24 h Clock Genes Expression in Human Adipose Tissue Explant Cultures. PLOS ONE. 2012;7(12):e50435. doi:10.1371/journal.pone.0050435

48. Petrus P, Mejhert N, Corrales P, et al. Transforming Growth Factor-β3 Regulates Adipocyte Number in Subcutaneous White Adipose Tissue. Cell Rep. 2018;25(3):551-560.e5. doi:10.1016/j.celrep.2018.09.069

49. Catalano-Iniesta L, Sánchez Robledo V, Iglesias-Osma MC, et al. Evidences for Expression and Location of ANGPTL8 in Human Adipose Tissue. J Clin Med. 2020;9(2):512. doi:10.3390/jcm9020512

50. Manzano-Núñez F, Arámbul-Anthony MJ, Albiñana AG, et al. Insulin resistance disrupts epithelial repair and niche-progenitor Fgf signaling during chronic liver injury. PLOS Biol. 2019;17(1):e2006972. doi:10.1371/journal.pbio.2006972

51. Cherubini A, Rusconi F, Lazzari L. Identification of the best housekeeping gene for RT-qPCR analysis of human pancreatic organoids. PLOS ONE. 2021;16(12):e0260902. doi:10.1371/journal.pone.0260902

52. López-Cerdán A, Andreu Z, Hidalgo MR, et al. Unveiling sex-based differences in Parkinson’s disease: a comprehensive meta-analysis of transcriptomic studies. Biol Sex Differ. 2022;13:68. doi:10.1186/s13293-022-00477-5

53. Català-Senent JF, Hidalgo MR, Berenguer M, et al. Hepatic steatosis and steatohepatitis: a functional meta-analysis of sex-based differences in transcriptomic studies. Biology of Sex Differences. 2021;12(1):29. doi:10.1186/s13293-021-00368-1

54. Pérez-Díez I, Hidalgo MR, Malmierca-Merlo P, et al. Functional Signatures in Non-Small-Cell Lung Cancer: A Systematic Review and Meta-Analysis of Sex-Based Differences in Transcriptomic Studies. Cancers. 2021;13(1):143. doi:10.3390/cancers13010143

55. Casanova Ferrer F, Pascual M, Hidalgo MR, Malmierca-Merlo P, Guerri C, García-García F. Unveiling Sex-Based Differences in the Effects of Alcohol Abuse: A Comprehensive Functional Meta-Analysis of Transcriptomic Studies. Genes (Basel). 2020;11(9):1106. doi:10.3390/genes11091106

56. Hall AN, Turner TN, Queitsch C. Thousands of high-quality sequencing samples fail to show meaningful correlation between 5S and 45S ribosomal DNA arrays in humans. Sci Rep. 2021;11(1):449. doi:10.1038/s41598-020-80049-y

57. Wang Z, Lachmann A, Ma’ayan A. Mining data and metadata from the gene expression omnibus. Biophys Rev. 2019;11(1):103–110. doi:10.1007/s12551-018-0490-8

58. The FAIR Guiding Principles for scientific data management and stewardship | Scientific Data. Accessed December 1, 2021. https://www.nature.com/articles/sdata201618

